# SAMWOOD: An automated method to measure wood cells along growth orientation

**DOI:** 10.64898/2026.03.11.711054

**Authors:** K. Verlingue, G. Brunel, A-L. Decombeix, M. Ramel, P. Tresson

## Abstract

Quantitative wood anatomy requires precise measurement of wood cells. This step is often laborious and limiting for further analysis. We introduce Samwood, a tool based on the zero-shot Segment-Anything Model to easily segment cells on microscopic images without the need for a training dataset. The reconstruction of cell files then allows for the analysis of wood along growth orientation and precise measurement of anatomical properties of the wood, such as lumen areas. We tested our pipeline on an example dataset of fossil woods featuring deformation, heterogeneous preservation, and frequent artefacts, to assess the robustness of our approach. The model achieves a precision of 0.78 and a recall of 0.80, often producing segmentation of better quality and more consistent than a human operator. This approach substantially reduces analysis time, minimizes operator bias, and provides a robust and extensible framework for large-scale anatomical studies The complete code pipeline is available at https://github.com/umr-amap/samwood.

**GRAPHICAL ABSTRACT:** 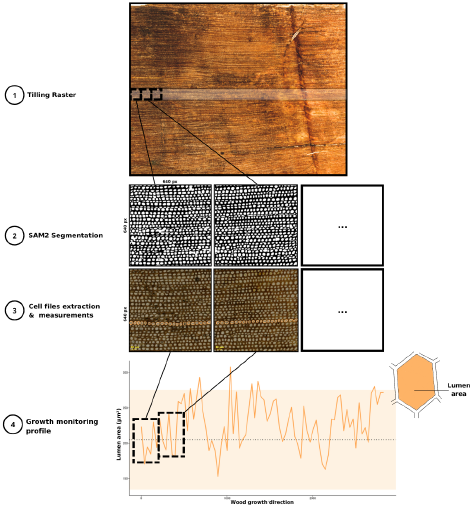

## 1 INTRODUCTION

Wood anatomy provides essential information for forest sciences, dendrochronology, and functional ecology, [2]. It also play a central role in understanding plant physiology and responses to environmental constraints [3]. Anatomical analysis of wood is based on microscopic observation of different cell types and their sizes. New cells are produced by the cambium during secondary growth, and experience variations driven by hormonal and environmental changes. In most cases, wood anatomical analyses of cells or growth rings, adopt a spatio-temporal perspective to track growth dynamics and tissue organization [4]. The production of large-area microscopic images of cross-sections enables the observation of the complete growth sequence, providing essential information for monitoring and understanding plant development [5].

However, the extraction of quantitative information from wood anatomical images still relies largely on time-consuming manual measurements [6, 7]. This represents a major methodological bottleneck in data acquisition, which restricts the amount of data that can be analyzed and limits the potential for large-scale studies[8].

Over the past decade, advances in machine learning and deep learning have transformed image analysis[9] and with it biological image analysis [10]. Deep learning models now enable the automated segmentation and measurement of complex biological structures, however, most deep learning applications in microscopy remain focused on human or medical cell imaging [11]. Few image analysis tools have been developed for plant tissues and cells [12, 13, 14], and even fewer are specifically adapted to the structural particularities of wood anatomy. Indeed, wood cellular structures are highly diverse and often discontinuous, yet consistently organized along growth gradients [15], which adds constraints for the image analysis of wood tissue.

To compensate the lack of large annotated datasets, benchmark and models, it is now possible to rely on foundation models. These models are trained on very large datasets and are afterwards able to generalize to a wide range of downstream tasks with little to no training.

One such model is SAM2, which is able to perform zeroshot object segmentation, *i*.*e*. to generate object masks without prior training on the target dataset or object categories[1]. In wood anatomy, large datasets are available but these are seldom annotated [16]. In this case, the use of zero-shot models like SAM2 seems appropriate to detect and measure wood cells.

Several studies have attempted to automate the extraction of quantitative wood-anatomical data in order to better understand endogenous mechanisms in trees, to clarify cellular organization, and to disentangle internal from external sources of variability. Early approaches focused on algorithmic detection of cell files, such as the method proposed by Brunel (2014) [5], which facilitated the extraction of cellular measurements from scanned images have been used to further automate image acquisition and cell segmentation, with WoodBot pipeline [3]. Other tools, such as ROXAS-AI[17], have also been developed to automatically identify tree-ring boundaries. However, these methods typically require model training, rely on specialized hardware, or remain limited in their generalizability.

Overall, the analysis of wood anatomical images comes with several constraints :

1. Measurements need to be accurate, but measuring cells has long been prohibitively time-consuming. The complexity of biological material and the already lengthy preparation required for microscopy make the process highly labor-intensive.
2. There is no large pre-existing and open annotated dataset.
3. Wood is structured and measurements need to be done following growth direction.

To answer these constraints, we present Samwood, an open-source Python package that automates the segmentation and measurement of woody plant cells from microscopic images of anatomical sections of wood. Samwood uses the SAM2 segmentation model to automatically outline cells without prior training. It first identifies cell rows and then extracts standardized quantitative measurements (area in µm^2^, width in µm, and double wall thickness in µm), significantly reducing analysis time and improving reproducibility.

## 2 MATERIAL AND METHODS

The analysis done in Samwood is performed in two majors steps: cell segmentation and cell files identification. Cell segmentation is done using SAM2 and integrating the constraints of working with large microscopic images. Cell file analysis is done by creating a cell connectivity graph. To illustrate the functioning of Samwood, we rely on an example dataset.

### 2.1 Example dataset

We compiled a dataset of 100 images (640 × 640 px) derived from digitalized thin sections of fossil wood and cellulose acetate peels, two preparation techniques commonly used to document anatomical structures in paleobotany [18, 19, 20]. The fossils material corresponds to specimens of Carboniferous, Permian and Triassic gymnospermous plant from various regions belonging to the collections of University of Kansas and University of Montpellier. A detailed list of specimen is provided as suplementaries informations. All the fossil woods were images using a Keyence VHX 7000 digital microscope (Keyence Corp., Osaka, Japan). A total of 8,980 manual annotations were generated using ImageJ to delineate individual tracheids i.e. the main type of cells in gymnospem wood that assure both mecanical support and water conductivity. These annotations provide a robust ground truth for evaluating segmentation performance. The full manual segmentation and numerization process represents an estimated 256 hours (2 months) of human work.

Fossil wood presents numerous artifacts due to the fossilization process and preparations methods. We therefore deemed that it was a good dataset to show the performances of the pipeline in a constraining context. The processing was also tested on extant wood (see Figure 5)

### 2.2 Segmentation of wood cells

Microscopic images are typically several orders of magnitude larger than the expected inputs of neural networks. Therefore, we have implemented a custom dataloader to extract square patches from raster images. When fed into the dataloader, an horizontal band of 640 pixel height and covering the entire width of the image is cropped in the original raster (see Figure 1). This band is then tiled into 640 × 640 pixel tiles to ensure complete coverage of the sample. The extracted patches are then processed using the segmentation model SAM2[1]. Prompts are given following a grid and objects are segmented by the model, with post processing steps handling double detections or noise. Several hyper-parameters can be tuned to influence the quality of segmentations such as the number of prompts, maximum size of object and allowed overlap. We tested a range of hyper-parameters combinations and selected a set that worked well to detect individual cells. To avoid false detections, objects above a given size threshold, deemed unlikely to correspond to biologically realistic values, are discarded. All cellular measurements are exported to a structured csv file containing cell identifiers, spatial coordinates, and quantitative measurements.

**FIGURE 1.**
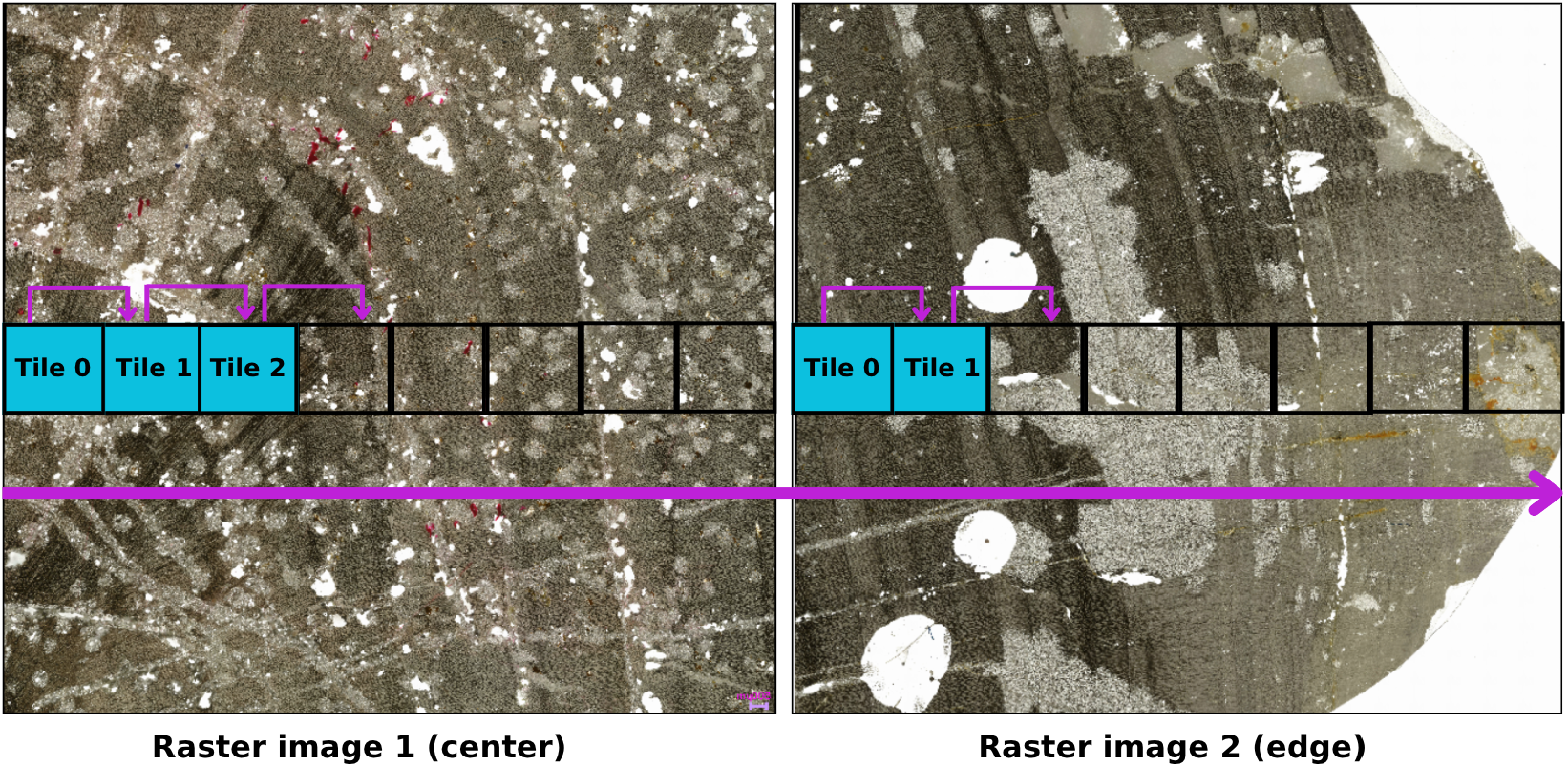
Example of tiling by the data loader on two images of fossil wood, one in the center and one near the outer edge.

**FIGURE 2.**
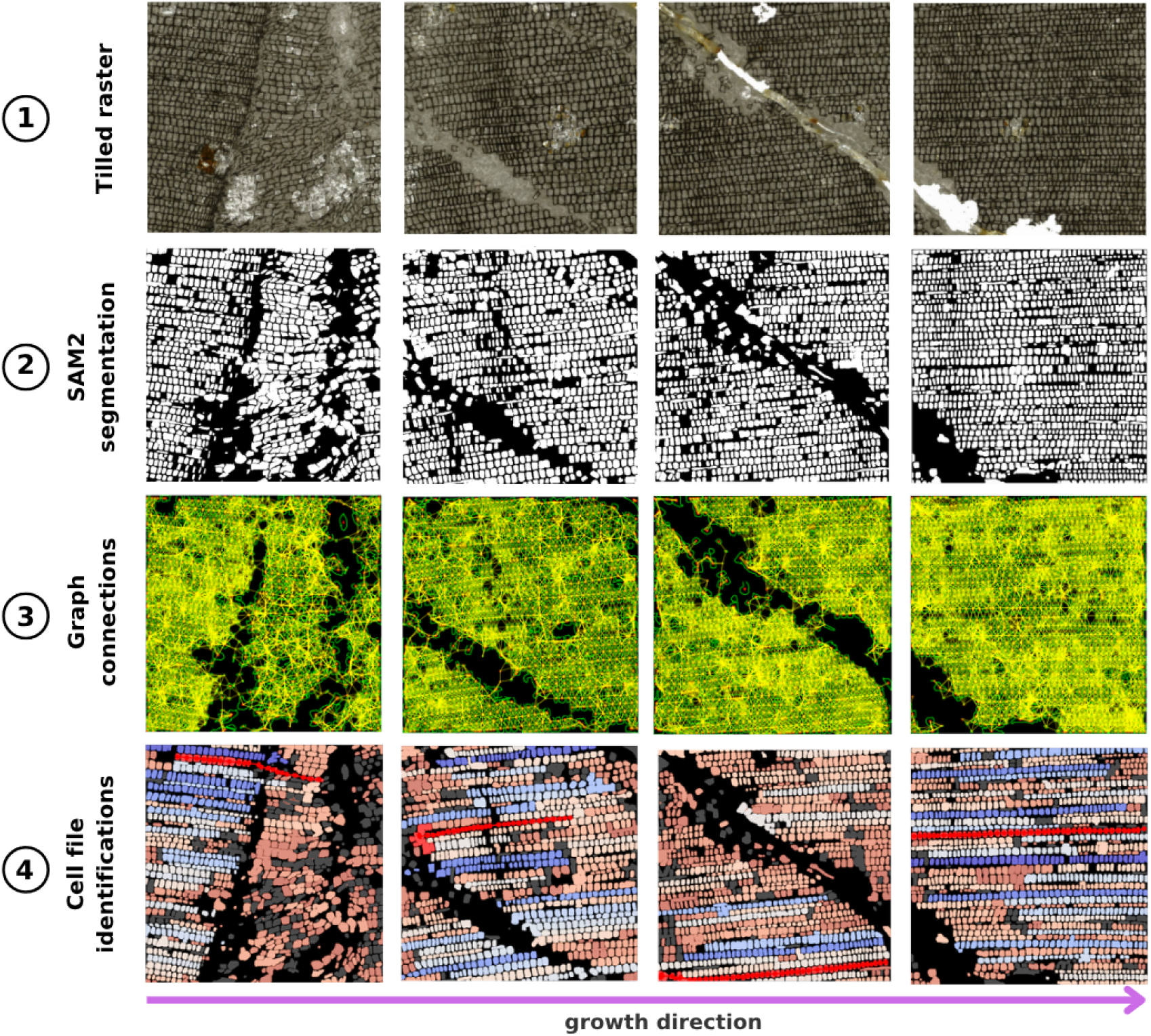
Overview of the processing. The original raster image is tilled (1), then tracheids are segmented using SAM2 (2). The masks are used to create the connectivity graph (3) and cell lines are identified (4).

### 2.3 Evaluation of segmentation

We evaluated the performance of SAM2 using our example dataset of fossil wood images manually annotated by human experts. Segmentations generated by SAM2 were compared with human labels using Intersection over Union, Recall, Precision and F1-score. To benchmark deep learning against lighter image-processing methods, we also performed colour-thresholding segmentation and compared the results with human annotations.

### 2.4 Cell files identification

Once tracheids were segmented, we developed an algorithm to identify and follow cell files across images following the method developed by [5]. Cell files correspond to linear series of cells derived from a single cambial initial over time [5], and their recognition is essential to track wood growth from the pith to the bark, particularly in deformed or poorly preserved fossil material. Cell boundaries are delineated using a watershed algorithm on the masks to separate adjacent cells. From these contours, a connectivity graph is constructed by linking cell centroids with edges. The angular distribution of edges is then analyzed to identify the dominant orientation, corresponding to the local wood growth direction. Cell files are traced iteratively by connecting nearest neighbors along this orientation, with a maximum distance threshold used to terminate a file when no valid neighbor is found. Each detected cell file is scored according to its length (number of cells), linearity, and morphological continuity (variation in cell area). For each tile, only the best-scoring file was retained.

### 2.5 Comparison with other methods

To evaluate the performance gain provided by the machine learning approach, we also performed tracheid segmentation using a traditional image thresholding method implemented in OpenCV 4.9.0. Segmentations obtained through this approach were compared both to human annotations and to those produced by the SAM2 model.

### 2.6 Target Metrics

For each identified cell file, individual cells are measured to quantify several morphological parameters. For each cell, the mask area, centroid coordinates (X, Y), and a unique cell identifier are recorded, allowing precise localization within the tile and the overall image. The equivalent diameter of each cell (*i*.*e*. assuming a circle) is calculated to provide a standardized measure of cell size.

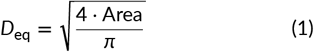

In addition, the thickness of the double cell wall is estimated by measuring the distance between centroids of adjacent cells, subtracting the portions overlapping the cell areas. This metric can only be reliably measured along the tracked cell files, where the connectivity of cells is preserved.

## 3 RESULTS

### 3.1 Segmentation accuracy

SAM2 produced segmentations closely matching human annotations (see Figure 3. Mean scores for Recall (0.80), Precision (0.78), and F1-score (0.79) indicate a good ability of the model to detect the target structures. However, the mean Intersection over Union (IoU = 0.68) highlights a lower overlap accuracy between the generated masks and manual annotations.

**FIGURE 3.**
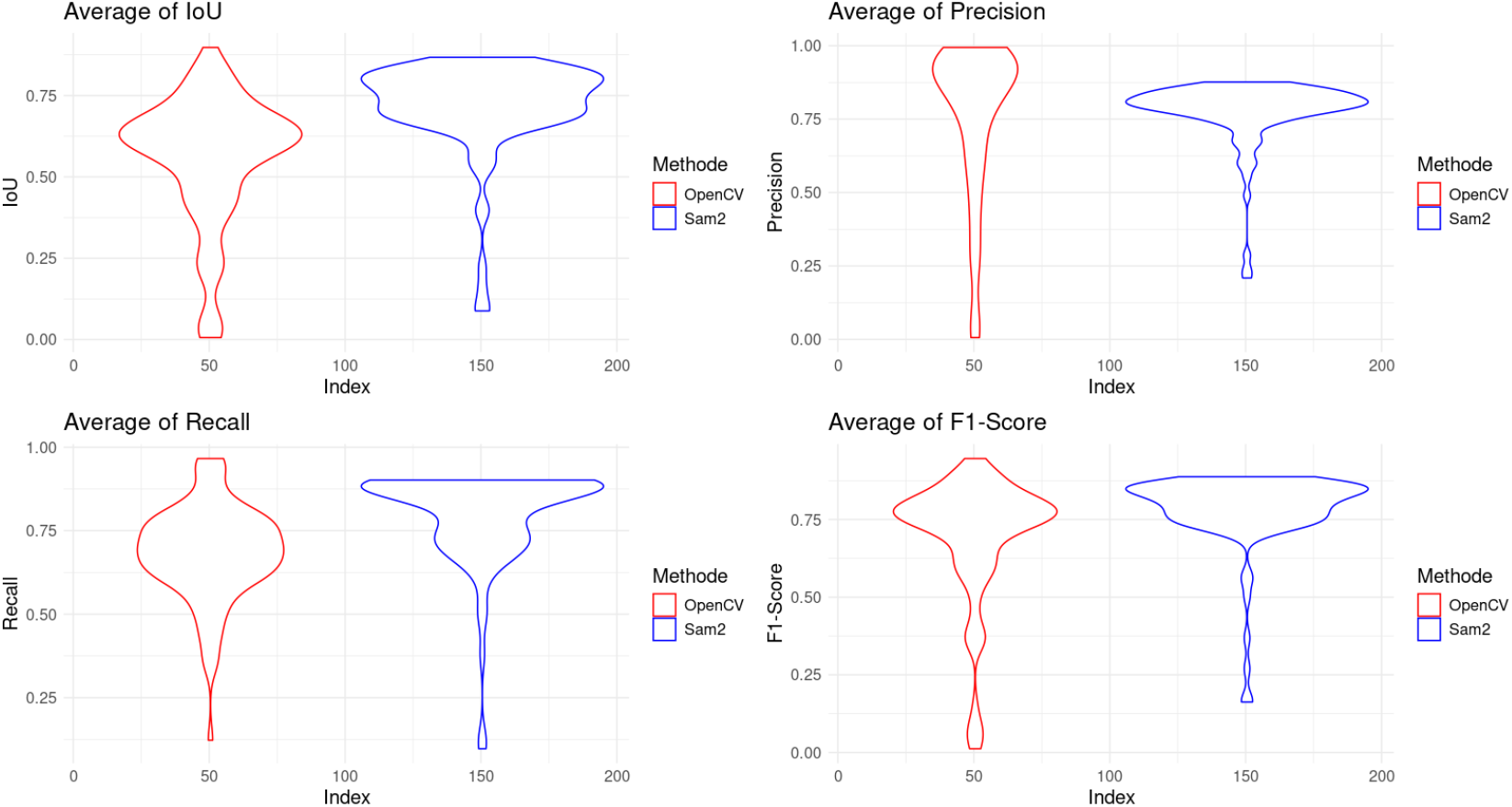
Comparisons of different evaluation metrics for the SAM2 segmentation model and a thresholding model relative to annotations made by a human expert.

### 3.2 Example analysis

## 4 DISCUSSION

### 4.1 Segmentation accuracy

As shown in figures 3 and 4, SAM2 is able to closely reproduce human labels without training. While recall and precision are high, IoU is lower, hinting at a shape difference in the segmentation produced by the model and those produced by humans. However, inspection of generated masks and human labels shows that generated masks may be of better quality than those produced by a human operator (see figure 4). The fact that we have worked on fossil wood is also comforting in the accuracy of the model, as samples are altered and provide more constraining conditions for the model than extant wood.

**FIGURE 4.**
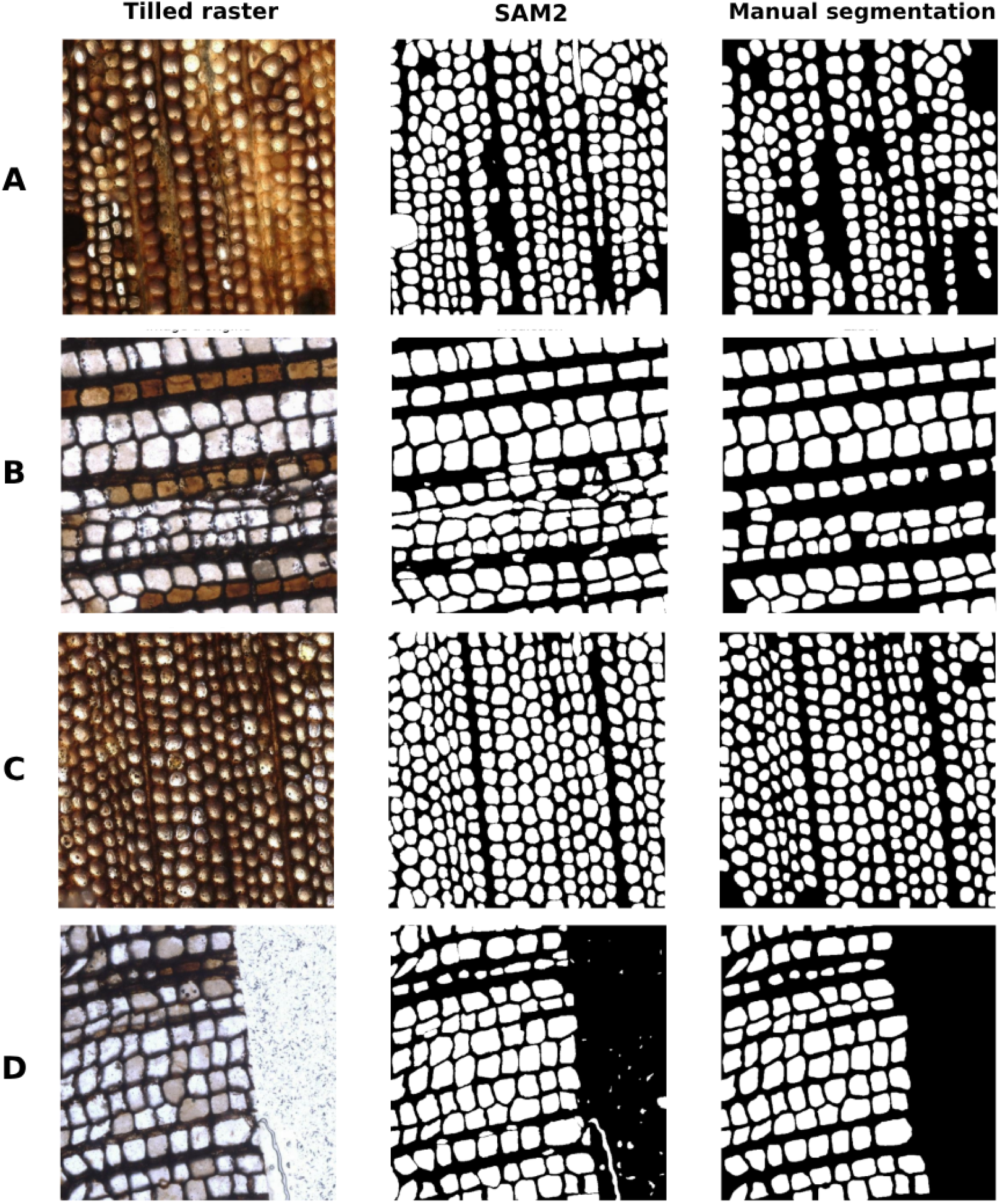
Example of tiles with human labels and SAM2 generated masks. The first and last examples show artifacts being segmented as objects. The second example shows the robustness of the model, even with obscured or colored cells. Close inspection shows that SAM2 segmentations are overall more detailed and more complete. Image tiled from specimens (A) ARX1XS1, (B) BOU1500AT, (C) ARX2XS1, (D) BOU1501DT

**FIGURE 5.**
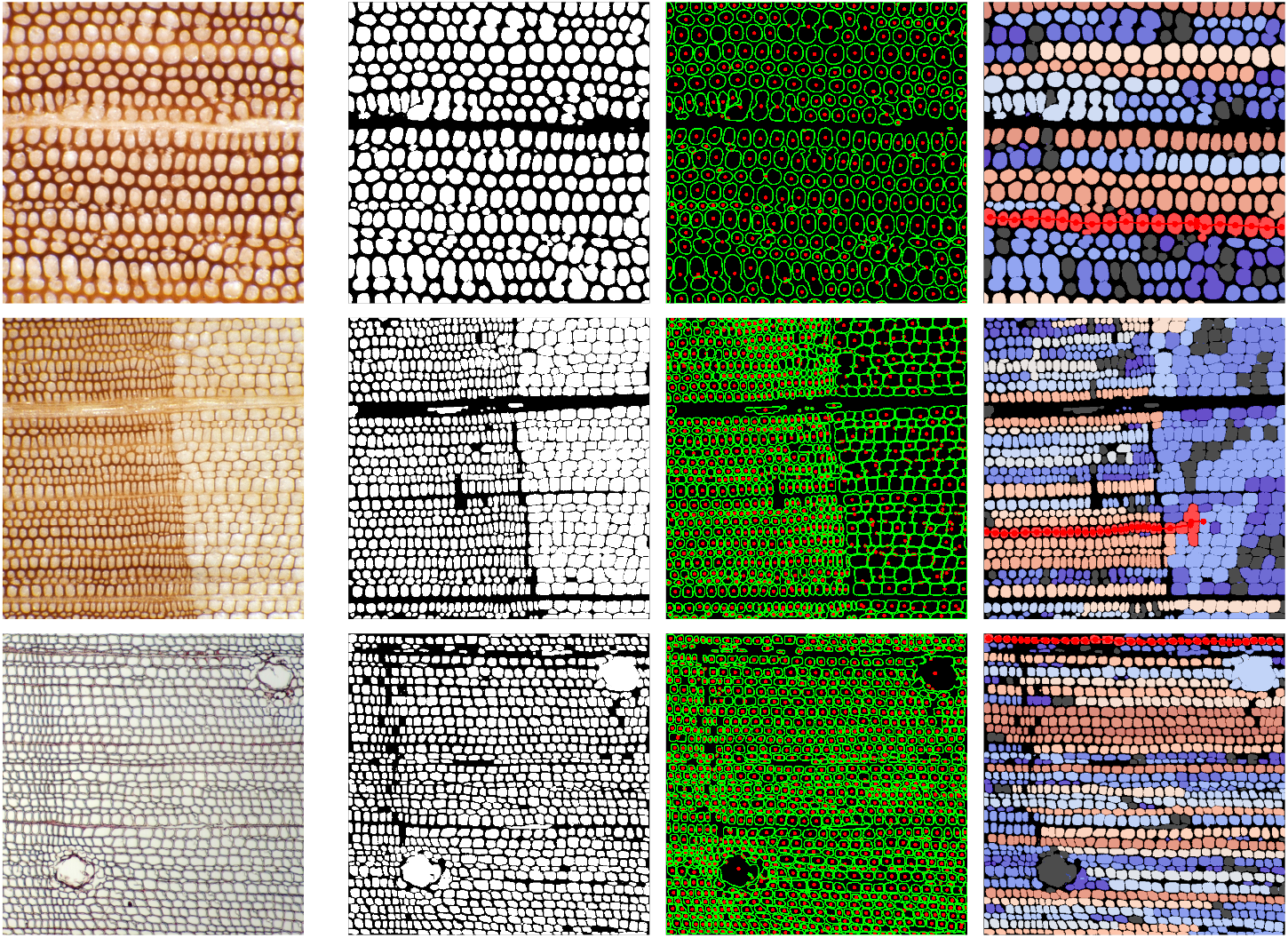
Tiles of modern wood samples and plots produced with our pipeline (masks, centroids, identified cell lines). From top to bottom, *Araucaria heterophylla, Picea orientalis* and *Pinus strobiformis*. These show the versatility of leveraging zero-shot detection. Images are taken from the InsideWood database: https://insidewood.lib.ncsu.edu/

### 4.2 Advantages of zero-shot

The zero-shot approach offers a major advantage for anatomical wood analysis, as it does not require any training data. This first step is indeed often the most important roadblock for leveraging automated image analysis as the creation of annotated training datasets is extremely costly in terms of time and expertise. In our case, the construction of the test set alone required more than 256 hours of manual work, to individually annotate nearly 8,980 cells. It would therefore be unrealistic to produce a training set of similar or larger size. Zero-shot segmentation bypasses this limitation by generating directly usable masks without a supervised training phase. Nevertheless, it requires minimal adjustment of a few hyperparameters and careful inspection of the generated masks, particularly to avoid false detections (artifacts) such as those illustrated in Figure 4, which tend to occur near the edges of thin sections (or when the preservation of fossil wood is heterogeneous or locally distorted).

Despite these issues, the method yields consistent performance and substantially reduces operator bias, which is a significant advantage for standardizing the extraction of anatomical descriptors. Ultimately, this ability to produce large numbers of measurements rapidly and in a reproducible manner opens the way for more precise monitoring of histological variation, as well as the creation of robust datasets for investigating growth patterns and environmental dynamics from wood cross-sections. Recently, SAM-3 has been released and may offer the possibility to incorporate example prompting [21], which in our context could enable even more precise segmentation by allowing users to explicitly indicate regions or objects that should not be segmented.

### 4.3 Versatility of the pipeline

The major strengths of our pipeline lies in its high versatility. Although it was originally designed to analyze fossil woods and track cellular files to extract functional traits relevant for (paleo)ecological applications, its architecture allows simple adaptation to a broad range of questions. In practice, the different components of the code (segmentation, cell files extraction and metric extraction) were built in a modular way, enabling rapid integration of new rules or additional analytical modules. As a result, the pipeline can be applied to tasks such as identifying growth-ring boundaries, measuring vessels in angiosperms, or detecting and excluding artefacts in altered or poorly preserved samples. The fact that it already performs well on fossil tissue characterized by deformation, heterogeneity and preservation suggests that it is sufficiently robust to be transferred to a wide variety of materials, including modern woods or images produced through different acquisition protocols (see. Figure 5). This versatility makes the pipeline a promising tool for standardizing and accelerating wood anatomical analysis across diverse research contexts.

## ACKNOWLEDGEMENTS

We would like to thank Brian Atkinson and Rudolph Serbet (Lawrence, KS, United States) for facilitating our access to fossils from the University of Kansas collections. We would also like to thank Stéphane Fourtier (AMAP) for his assistance during the development process. Finally, we would like to thank AMAP (botAny and Modelling of Plant Architecture and vegetation), a joint research unit bringing together the University of Montpellier, the CNRS (UMR5120), CIRAD (UMR51), INRAe (UMR931) and IRD (UR123).

## 5 SUPPORTING INFORMATION

List of fossils informations in Table 1.

**TABLE 1.**
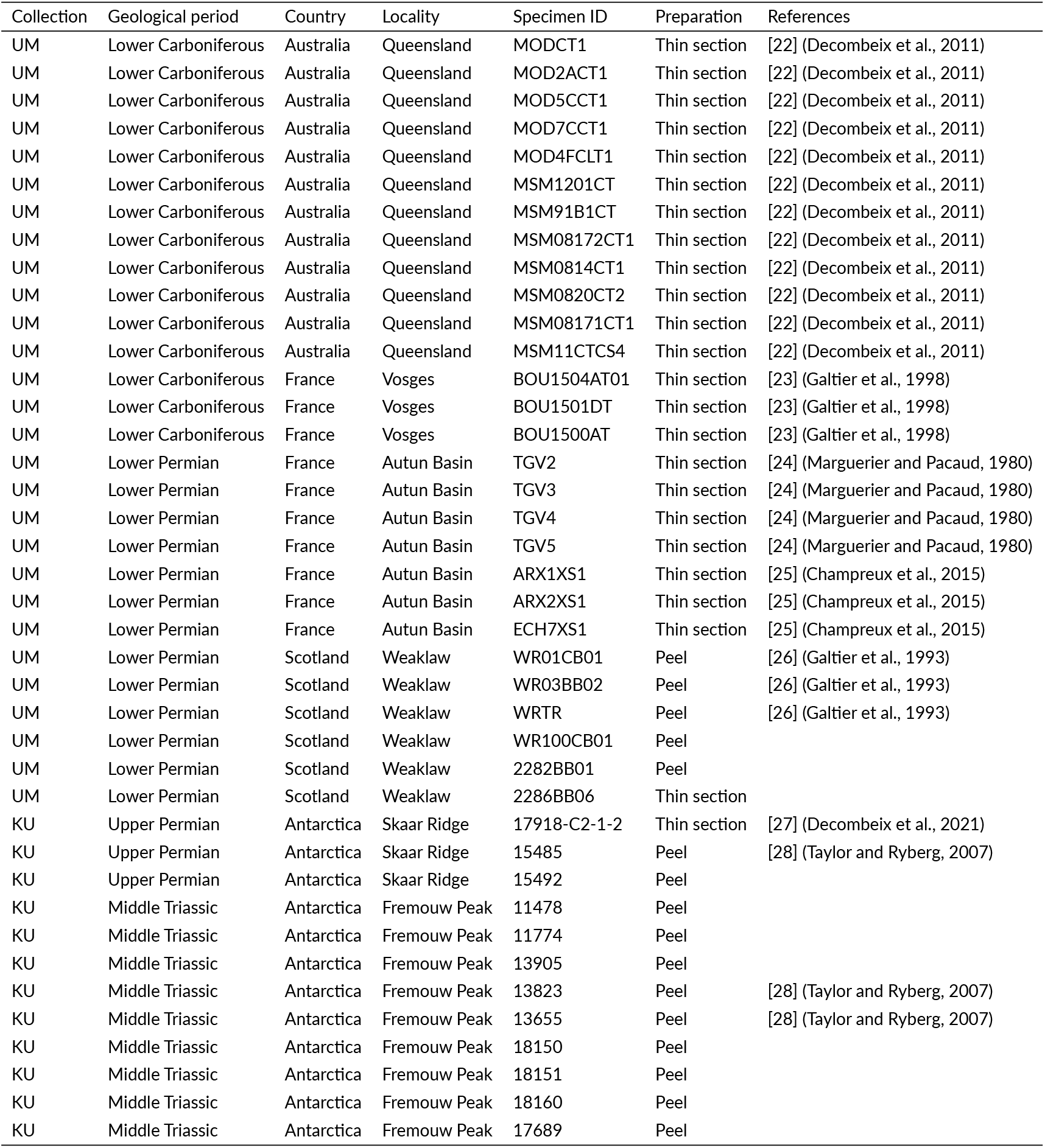
Summary of fossil wood specimens used in this study.

Visual comparison of 100 visual comparisons segmentations between SAM 2 and manual segmentation : https://doi.org/10.6084/m9.figshare.29136194. v1.

Numerised images of fossilised wood from the University of Montpellier (UM) collection : https://figshare.com/s/9e9470dd445909be0338

